# Phase-based genetic logic circuits

**DOI:** 10.1101/2022.12.13.520289

**Authors:** Timothy J. Rudge, Gonzalo Vidal

**Affiliations:** Interdisciplinary Computing and Complex Biosystems, School of Computing, Newcastle University, Newcastle upon Tyne

**Keywords:** Genetic circuit, logic, noise, phase

## Abstract

Bio-computation is the implementation of computational operations using biological substrates, such as cells engineered with synthetic genetic circuits. These genetic circuits can be composed of DNA parts with specific functions, such as promoters that initiate transcription, ribosome binding sites that initiate translation, and coding sequences that are translated into proteins. Compositions of such parts can encode genetic circuits in which the proteins produced by genes regulate each other in different ways. The expression level of each gene may be considered as a signal which may be high or low, encoding binary logic, and the combinations of genes can then encode logic circuits. This is equivalent to the level-based logic used in modern electronic computers in which the voltage forms the logical high or low states.

A different approach to binary logic is to encode the high and low states in the phase of an oscillating signal. In this approach a signal in phase with a reference represents high, and a signal antiphase with the reference represents the logical low state. We present here designs and models for phase-based genetic NOT, OR, AND, MAJORITY and complementary MAJORITY gates, which together form multiple complete logical sets. We derive analytical expressions for the optimal model parameters for circuit function. Our simulation results suggest that this approach to genetic logic is feasible and could be less sensitive to gate input-output mismatch than level-based genetic logic. To demonstrate the scaleability of phase-based genetic logic, we used our complementary MAJORITY and NOT gates to design and simulate a fully functional 4-bit ripple adder circuit, and showed that it was robust to molecular noise.

## 1 Introduction

From the earliest conceptual models genes have been considered as Boolean computational elements, which may be expressed or not, representing logical 0 and 1 states (*1*). This concept has been exploited by Synthetic Biologists to construct genetic circuits, analogous to their electronic counterparts in which high and low voltages represent the logical states. This is level-based binary logic. Biocomputation can be performed using these circuits, providing a means to encode defined information processing in living cells (*2*). For example genetic NOT gates can be implemented using genes containing repressible promoters such that when the input repressor is present the output gene expression is low (logical 0) and vice versa (*3*). Extending this principle, tandem repressible promoters can encode NOR gates (*4*), which are functionally complete - that is, they can be combined to produce any logic circuit (*5*). Other approaches using riboswitches have created NAND gates which are also functionally complete (*6*). Although this approach is powerful it suffers from several drawbacks. Firstly it is sensitive to noise inherent in gene expression. Also, it is sensitive to input-output matching between gates, which requires accurate estimation of gate parameters to enable reliable design (*5*). These effects can combine to produce logical glitches and hazards (*7*). Finally, it is not necessarily efficient in terms of the number of gates required to construct a particular logic circuit.

This presents a need for alternative or non-conventional approaches to biocomputation. One such approach is phase-based binary computation. Phase-based binary logic encodes the high and low states in the phase of an oscillating signal. In this way a signal in phase with a reference represents logical 1, and a signal antiphase with the reference represents the logical 0 state. John von Neumann (*8*) and Eichi Goto (*9*) both developed the principles of phase-based computation using electrical oscillators and demonstrated that MAJORITY, NOT, AND, and OR gates can be made by combining oscillating signals. Phase-based logic has been shown to be less sensitive to noise than level-based logic (*10*). The ease of construction of a range of logic gates, including the MAJORITY operation may also allow more efficient designs for some circuits. Although these ideas are known in computing, to the best of our knowledge they have not been implement in biological systems. We present here designs and models for phase-based genetic logic gates, derive analytical expressions for their optimal parameters, and use simulations to explore the feasibility of constructing circuits from them.

## Results

Similar to the level-based logic paradigm, gene expression levels form the basis of our logical states, whereas in our case these levels will oscillate. In order to model the oscillating gene expression levels, we used a one-step protein synthesis model. Such a model can be formulated using the stochastic simulation algorithm (*11*) with protein production and degradation reactions, such that any genetic circuit may be modeled as follows,

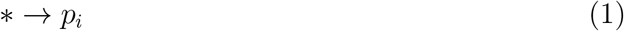

with propensity *ϕ*_*i*_(*s*_0_, *s*_1_…*s*_*m*_, *p*_0_, *p*_1_…*p*_*n*_), and

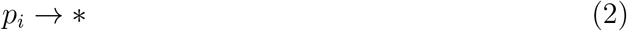

with unit propensity, normalizing time by the protein degradation time scale. Here the *p* are the *n* proteins produced by each gene in the circuit, and *s* are the *m* input signals. The functions *ϕ*_*i*_ encode the interactions between gene *i* and the proteins produced by the circuit and signal molecules added to the cell culture. The steady-state protein level is given by *ϕ*_*i*_(*s*_0_, *s*_1_…*s*_*m*_, *p*_0_, *p*_1_…*p*_*n*_).

### Input signal receivers

The inputs to our logic circuits come from typical one-component chemical receiver circuits (*12*), thus the output is driven by a single inducible promoter (figure 1). Assuming that the receptor protein is at saturating levels, we may model these signal receivers using the simple Hill function,

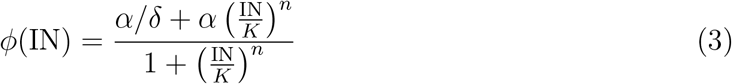

where IN is the input signal concentration, *K* is the switching point, *n* is the cooperativity, *α* is maximal expression rate, and *δ* is the dynamic range.

**Figure 1:**
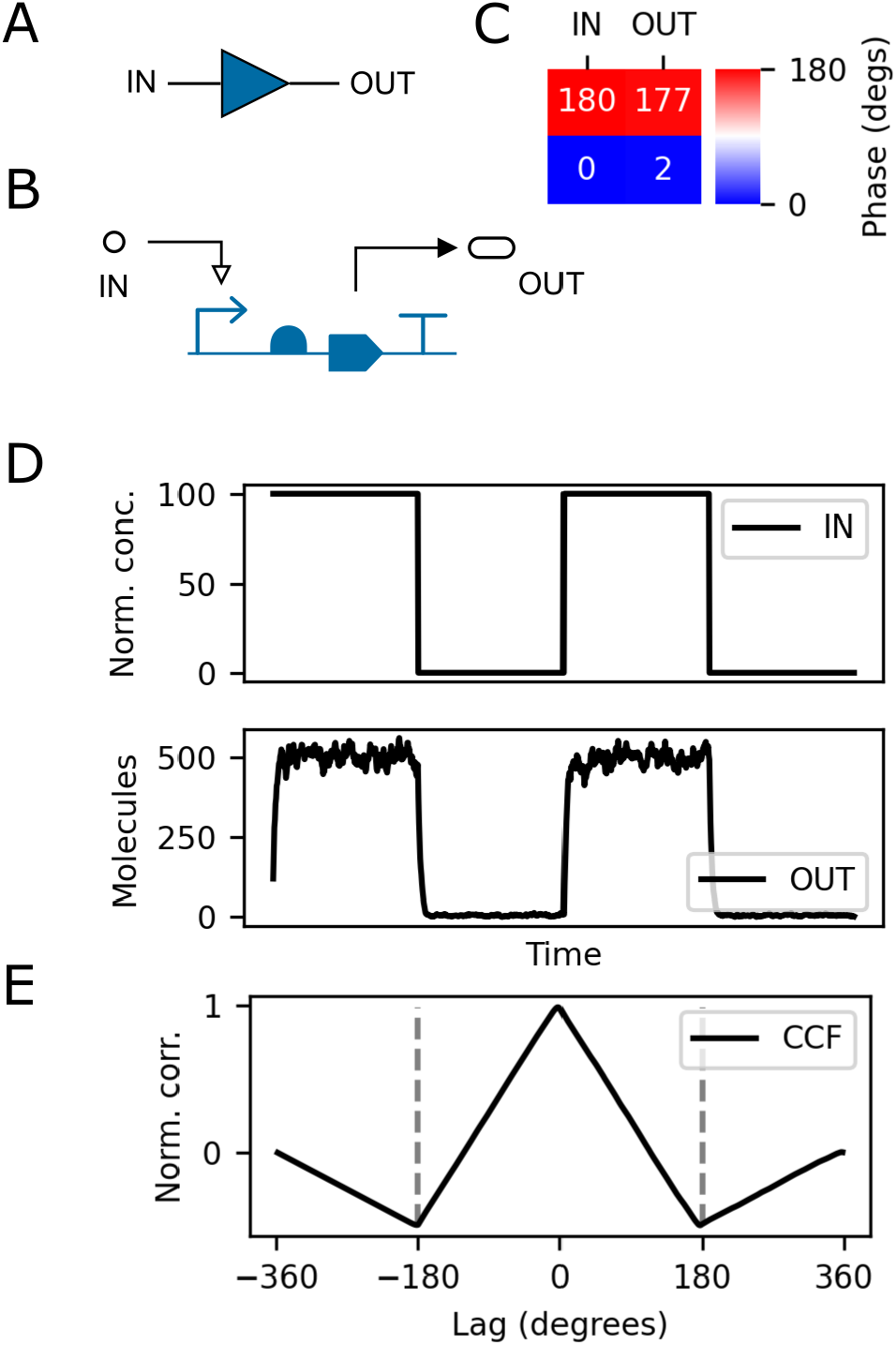
Design of input signal receiver. (A) Symbolic representation showing input and output. (B) Genetic implementation using an inducible promoter. (C) Logical truth table showing mean phase of 100 stochastic simulations, where antiphase indicates logical 0 and in phase represents logical 1. (D) Example stochastic simulation traces for normalized input signal concentration and output protein level. (E) Normalized cross-correlation function (CCF) showing a peak at 0 degrees indicating that the output is in phase with the input.

We envisage that these signals are provided to the cells at oscillating concentrations with some fixed period and phase. The most practical method of doing this would involve switching media with and without inducer chemical, such that the input signal would be a square wave. We choose one of these signals as the reference and set the other signals either in phase (logical 1) or antiphase (logical 0).

To produce a square wave we would like to drive the signal receiver between its minimal and maximal expression rates. However we are limted by the maximum signal concentration that can be tolerated by the cell, and the leaky expression rate in the absence of signal. A reasonable range of signal inputs would be between from zero to two orders of magnitude above the switching point *K*. Let us define the minimal and maximal output protein levels as *p*_*OFF*_ and *p*_*ON*_. Then we find that *p*_*OFF*_ = *α/δ* and assuming *n* is such that 1*/δ* ≪ 10^2*n*^,

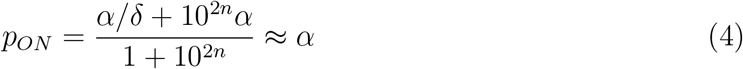

Figure 1D shows example stochastic simulations with *α* = 500, and figure 1E shows the resulting normalized cross-correlation function (CCF). The CCF displays a clear peak at 0 degrees indicating that the output is in phase with the input as required. We performed 100 stochastic simulations with *α* = 500 confirming that this phase relationship was maintained despite molecular noise.

We now wish to choose the switching point *K*_*D*_ for the downstream gates such that it is centred on the output dynamic range of the receiver in log space. This means that *K*_*D*_ is the geometric mean of the on and off protein levels,

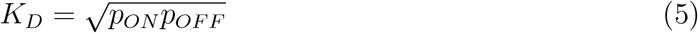

and so 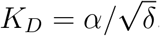. The output dynamic range of the receiver is *δ*_*OUT*_ = *p*_*ON*_ */p*_*OFF*_ = *δ*.

### NOT gate

NOT gates can be formed by simply inverting the input signal, which can be achieved if the protein *p* is a repressor corresponding to a downstream promoter (figure 2). This design is identical to a typical level-based NOT gate (*3*). In this case, when the input signal is high, the promoter output is low and vice versa, so that the output signal is antiphase with respect to the input and thus is its logical inverse in phase representation. The model is,

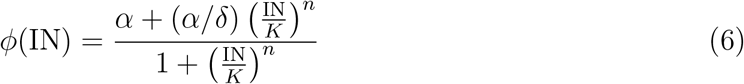

where IN is the input protein level. Setting *K* to the optimal defined above, *α* = 500 and *δ* = 100, we simulated a circuit consisting of an input signal receiver connected to a NOT gate (figure 2). The results show that the antiphase relation was maintained despite the intrinsic molecular noise.

**Figure 2:**
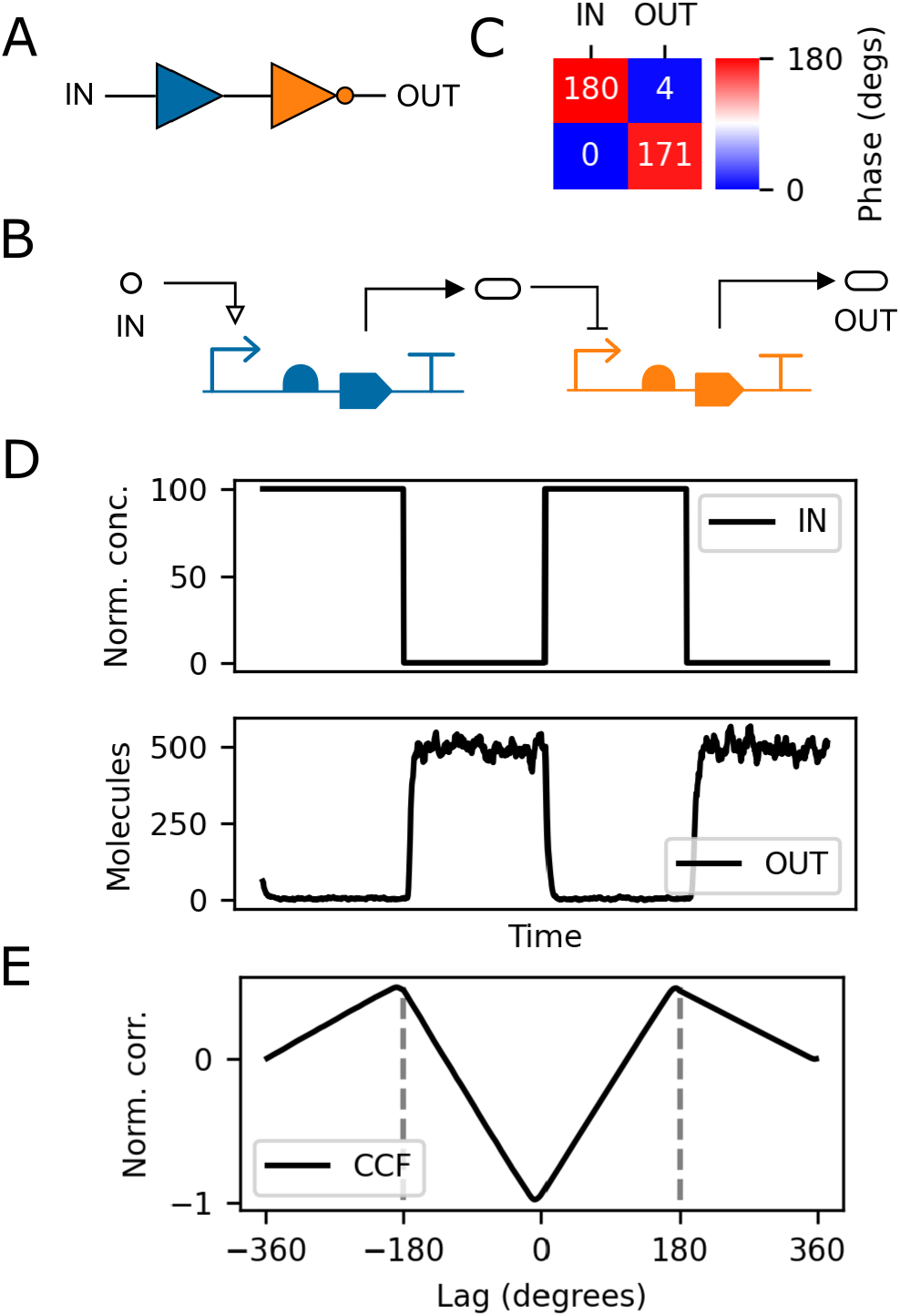
Design of phase-based NOT gate. (A) Schematic showing an input signal receiver connected to a downstream NOT gate. (B) Genetic implementation of the circuit described in A. (C) Logical truth table showing mean phase of 100 stochastic simulations, where antiphase indicates logical 0 and in phase represents logical 1. (D) Example stochastic simulation traces for input normalized signal concentration and output protein level showing antiphase relation. (E) Normalized cross-correlation function (CCF) showing peak at approximately 180 degrees, indicating that the input and output are antiphase as required.

The input has its own dynamic range *δ*_IN_, and is centered on the switching point *K* in log space so that we have,

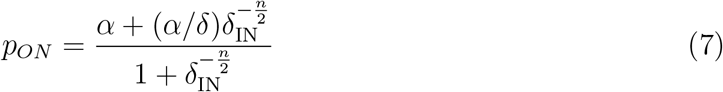

and,

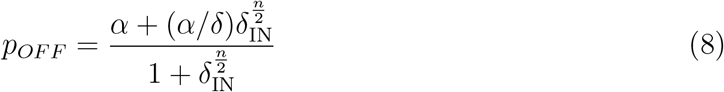

To center the output of the NOT gate to match the downstream gate, we choose the downstream switching point *K*_*D*_ such that,

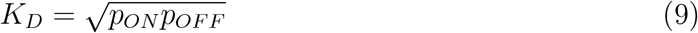

and so,

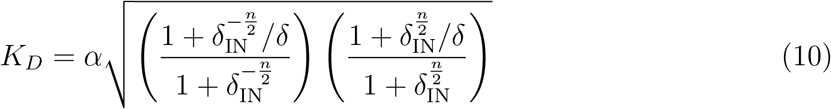

The actual output dynamic range of the NOT gate given input dynamic range *δ*_IN_ is,

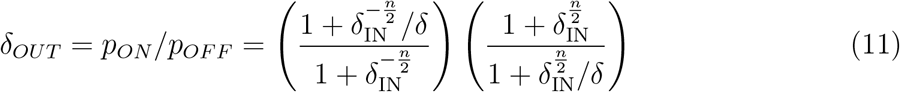

so that as *δ*_IN_ → ∞, *δ*_*OUT*_ → *δ*.

### Gate input-output mismatch

We have derived the optimal switching point for gates downstream of the signal receiver and NOT gate. However, in practice available parts will not exactly match these parameters. Furthermore, experimental estimates of gate parameters are subject to uncertainty due to, for example, their context in the application genetic system. This may lead to circuit failure, especially in circuits in which signals must pass through multiple gates before generating the output. In order to test this we simulated chains of NOT gates in which the output of one gate serves as the input to another downstream NOT gate (figure 3A). We considered chains of 1, 3, 5, 7 and 9 gates and performed 100 stochastic simulations of both level-based and phase-based inputs. Maximal steady state protein levels *α* were chosen as 500 molecules, giving an ideal switching point of approximately 50 molecules, consistent with estimates for the LacI repressor (*13*). For each simulation each gate switching point *K* was subject to mismatch such that,

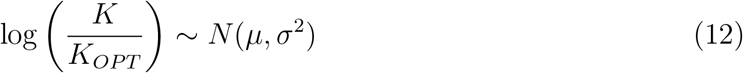

where *N* (*μ, σ*^2^) is a normal distribution with mean *μ* and standard deviation *σ*, and *K*_*OPT*_ is the calculated optimal switching point as derived above. We used *μ* = 0 and 2*σ* = log(10), meaning that the mismatch ranges over approximately one order of magnitude below and above the ideal value (figure 3B). We then quantified the error rate defined as the proportion of simulations that gave incorrect final output. The phase-based approach gave consistently lower error rate for all circuit sizes (figure 3C).

**Figure 3:**
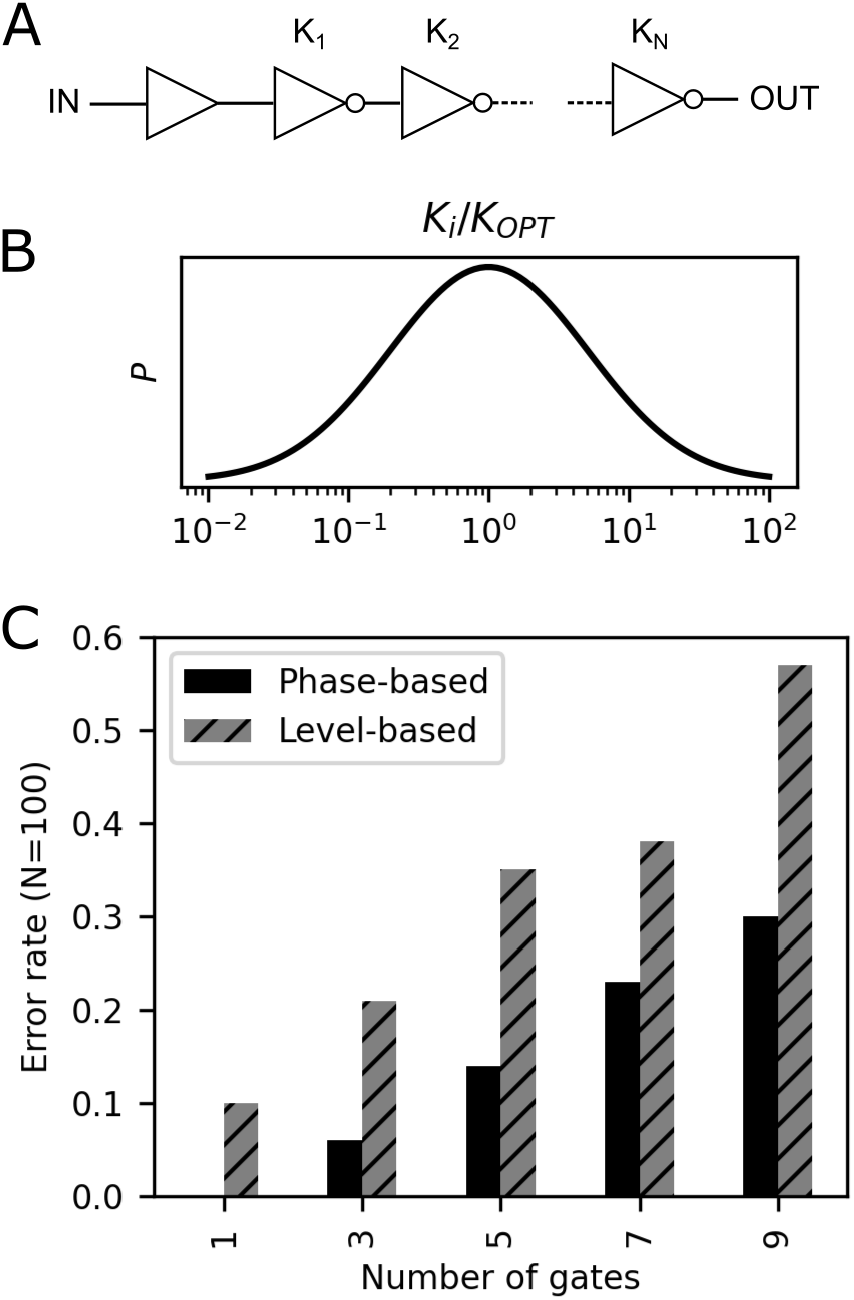
Effect of gate mismatch on circuit performance. (A) NOT gates were connected to form chains in which the output of one gate serves as the input to the next. (B) The switching points of the gates were set randomly according to a log normal distribution, where *K*_*OPT*_ is the analytically derived optimal value. (C) The error rate from 100 stochastic simulations for chains of different lengths shows that phase-based logic outperforms level-based logic.

### MAJORITY gate

A nice feature of oscillating signals is that we can exploit constructive and destructive interference to perform computation. When signals are summed they will interfere. Signals can be summed by using multiple promoters in series, such that the output of the gene is the sum of the outputs of each promoter. Consider a gene driven by three inducible promoters. When two or more of the signals are in phase, the resulting output is in phase, and vice versa. When we consider the phase of the protein levels as logical states, this is the MAJORITY operation. We model the gate as follows,

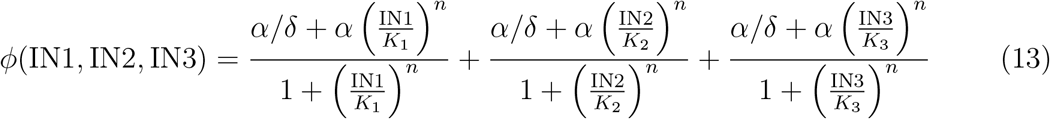

where IN1, IN2, and IN3 are the three input activator protein levels. Using the optimal switching level derived for the signal receiver above, we simulated a MAJORITY gate (*α* = 500, *δ* = 100) connected to three input signal receivers (figure 4). The resulting phase relationships between input and output of 100 stochastic simulations gave the correct logical truth table (figure 4C).

**Figure 4:**
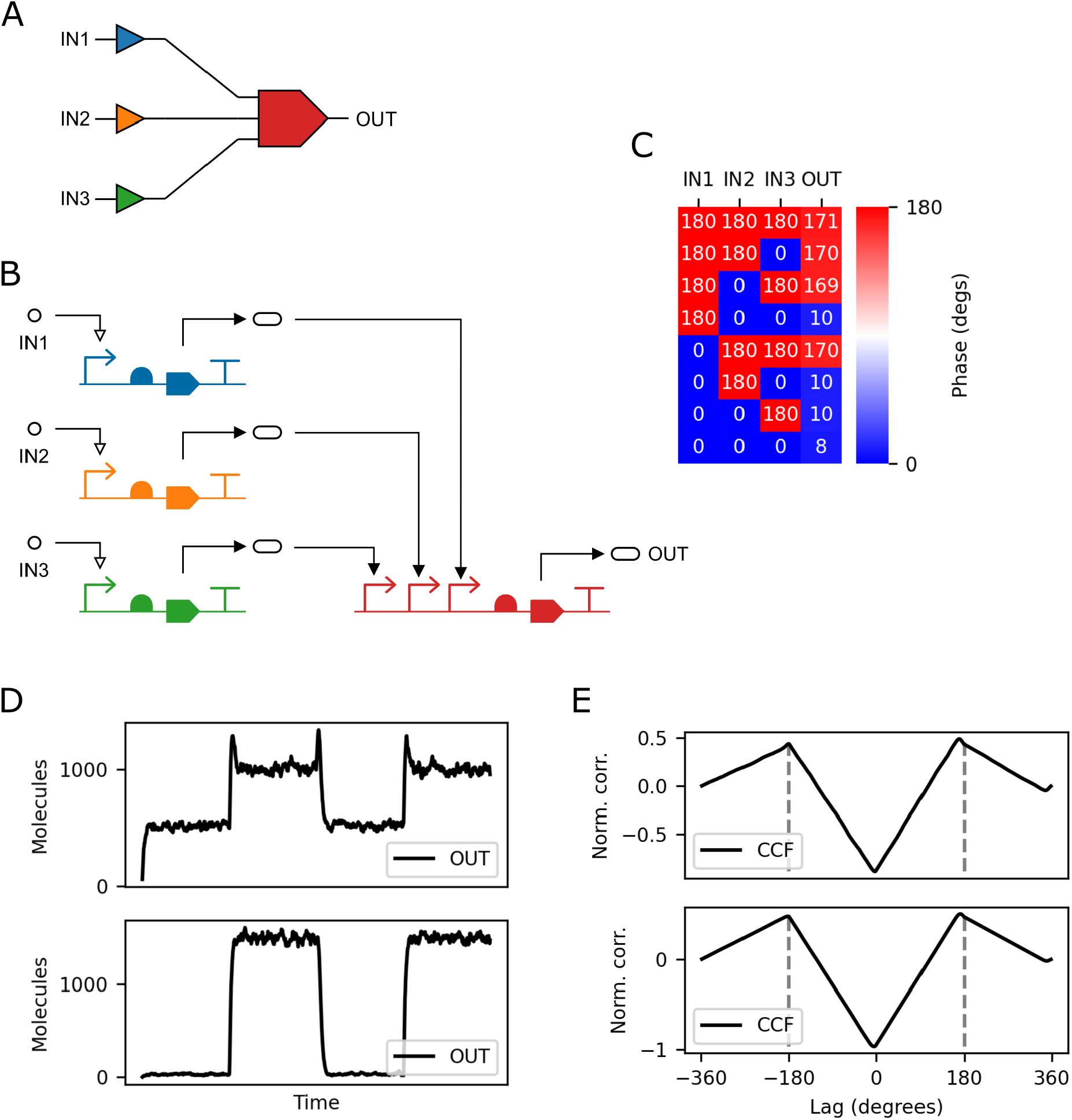
Design of phase-based genetic MAJORITY gate. (A) Schematic of a simple test circuit with three input signal receivers connected to a downstream MAJORITY gate. (B) Genetic implementation of the circuit in A. (C) Logical truth table showing mean phase of 100 stochastic simulations, where antiphase indicates logical 0 and in phase represents logical (D) Example stochastic simulation traces for input normalized signal concentration and output protein level for worst (top) and best (bottom) possible output dynamic ranges. (E) Normalized cross-correlation functions with respect to the reference signal for the examples in D.

For simplicity let us assume that the inputs have equal dynamic range *δ*_IN_, so the IN1*/K*_1_, IN2*/K*_2_, IN3*/K*_3_ range from 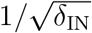 to 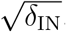. The worst case output dynamic range is when two inputs are in phase, then we have, assuming that all input dynamic ranges are equal,

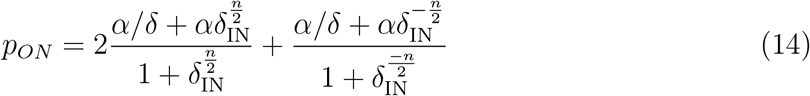

and,

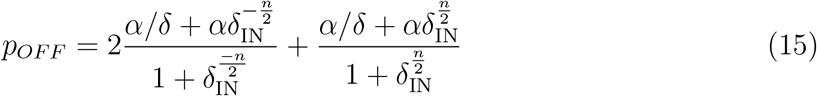

This leads to the optimal *K*_*D*_ for downstream gates,

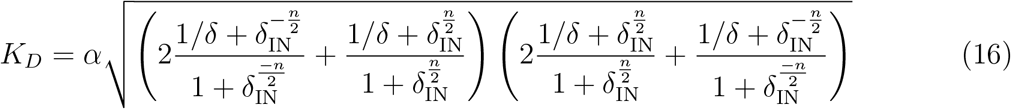

The worst case actual dynamic range of the output of the gate is *δ*_*OUT*_ = *p*_*ON*_ */p*_*OFF*_, which when *δ*_IN_ is large tends to,

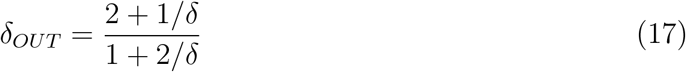

The maximal achievable output dynamic range is therefore *δ*_*OUT*_ = 2. This limited output dynamic range can be seen in the example stochastic simulation traces shown in figure 4D. This analysis is easily extended to the case where each input has different dynamic range.

### Complementary MAJORITY gate

The MAJORITY gate described above has limited dynamic range compared with the NOT gate. This means that signals would rapidly degrade as they pass through circuits. In order to amplify the dynamic range we can pass the output to a NOT gate, inverting the signal and forming a complementary MAJORITY gate. To see this recall the output dynamic range of the NOT gate as a function of its input dynamic range (equation 11), and consider the case as the dynamic range of the NOT gate *δ*_*NOT*_ → ∞,

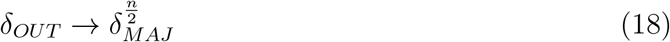

where *δ*_*MAJ*_ is the output dynamic range of the MAJORITY gate as defined above. Hence if *n >* 2 the output dynamic range of the MAJORITY gate is effectively amplified. In this study we use *n* = 4, as is the case of the LacI repressor protein (*14*), so that the dynamic range is squared after passing through the NOT gate.

### NOR and NAND gates

Using the MAJORITY gate described above, combined with a reference input signal it is possible to perform OR and AND operations. By feeding the reference signal into a receiver that then produces an activator of the MAJORITY gate, say input IN3, we see that the result is an OR gate. When either of the signals IN1 or IN2 is in phase with the reference IN3 the result of the sum is in phase with the reference. Conversely if the input IN3 is set antiphase with respect to the reference, we form the AND operation. When both IN1 and IN2 are in phase with the reference they are antiphase with IN3 and so the result is antiphase. In all other cases the result is in phase with the reference. Again the output dynamic range of these gates is limited, and we can amplify it by passing through a NOT gate to form NOR and NAND gates. The same procedure as applied for the MAJORITY gate described above can be used to derive optimal parameters for input-output matching.

### Phase-based arithmetic

Next we designed a phase-based full adder genetic circuit using only complementary MAJORITY and NOT gates (figure 6A,B). The circuit consists of three complementary MAJORITY gates and two NOT gates. Each gate is connected to downstream gates via transcription factors (activators or repressors). In this design we used an operon producing both a repressor and an activator from the same transcript by including two RBS (ribosome binding sites) and two CDS (coding DNA sequence).

**Figure 5:**
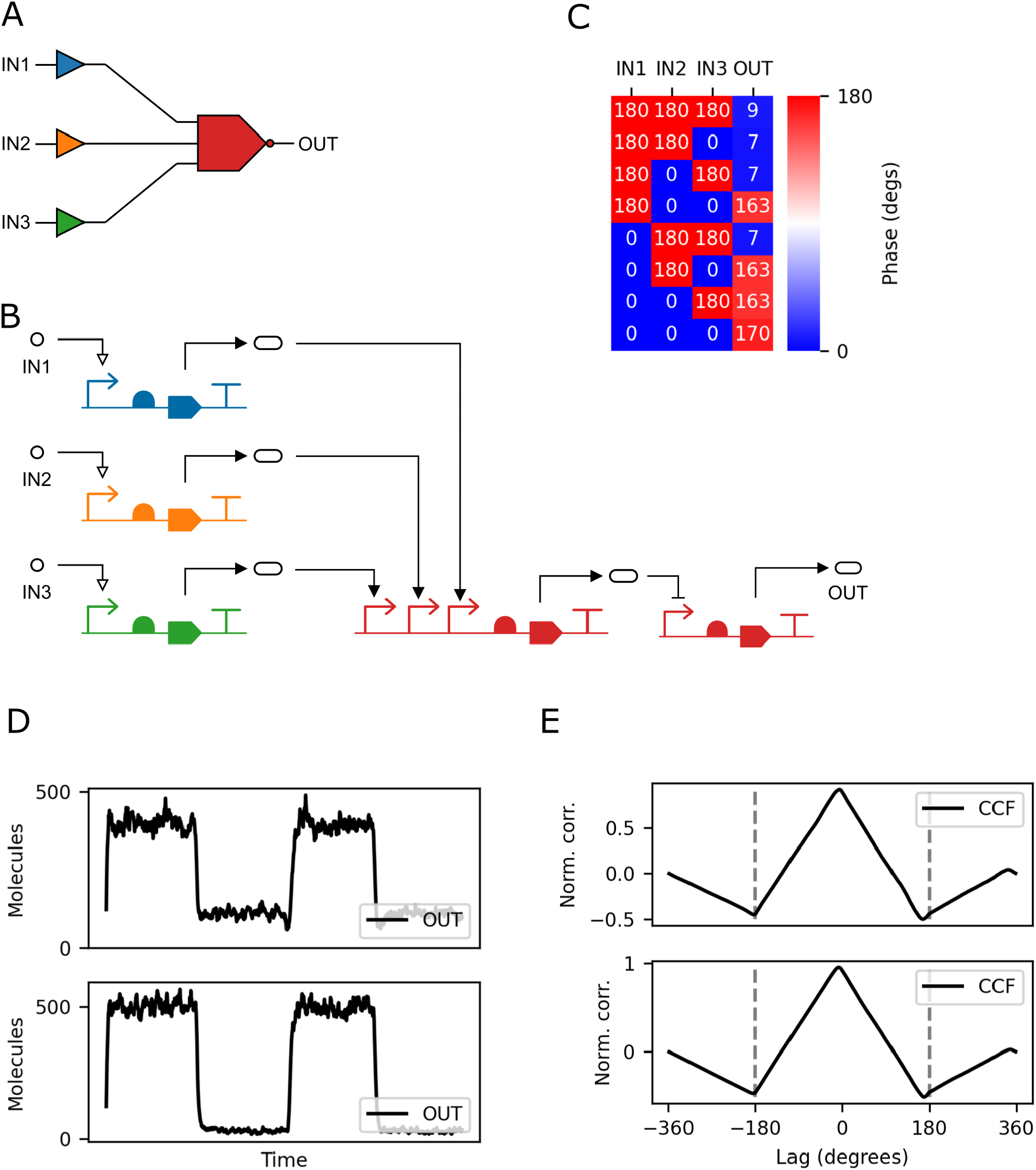
Design of phase-based genetic complementary MAJORITY gate. (A) Schematic of a simple test circuit with three input signal receivers connected to a downstream complementary MAJORITY gate. (B) Genetic implementation of the circuit in A. (C) Logical truth table showing mean phase of 100 stochastic simulations, where antiphase indicates logical 0 and in phase represents logical 1. (D) Example stochastic simulation traces for input normalized signal concentration and output protein level for worst (top) and best (bottom) possible output dynamic ranges. (E) Normalized cross-correlation functions with respect to the reference signal for the examples in D.

**Figure 6:**
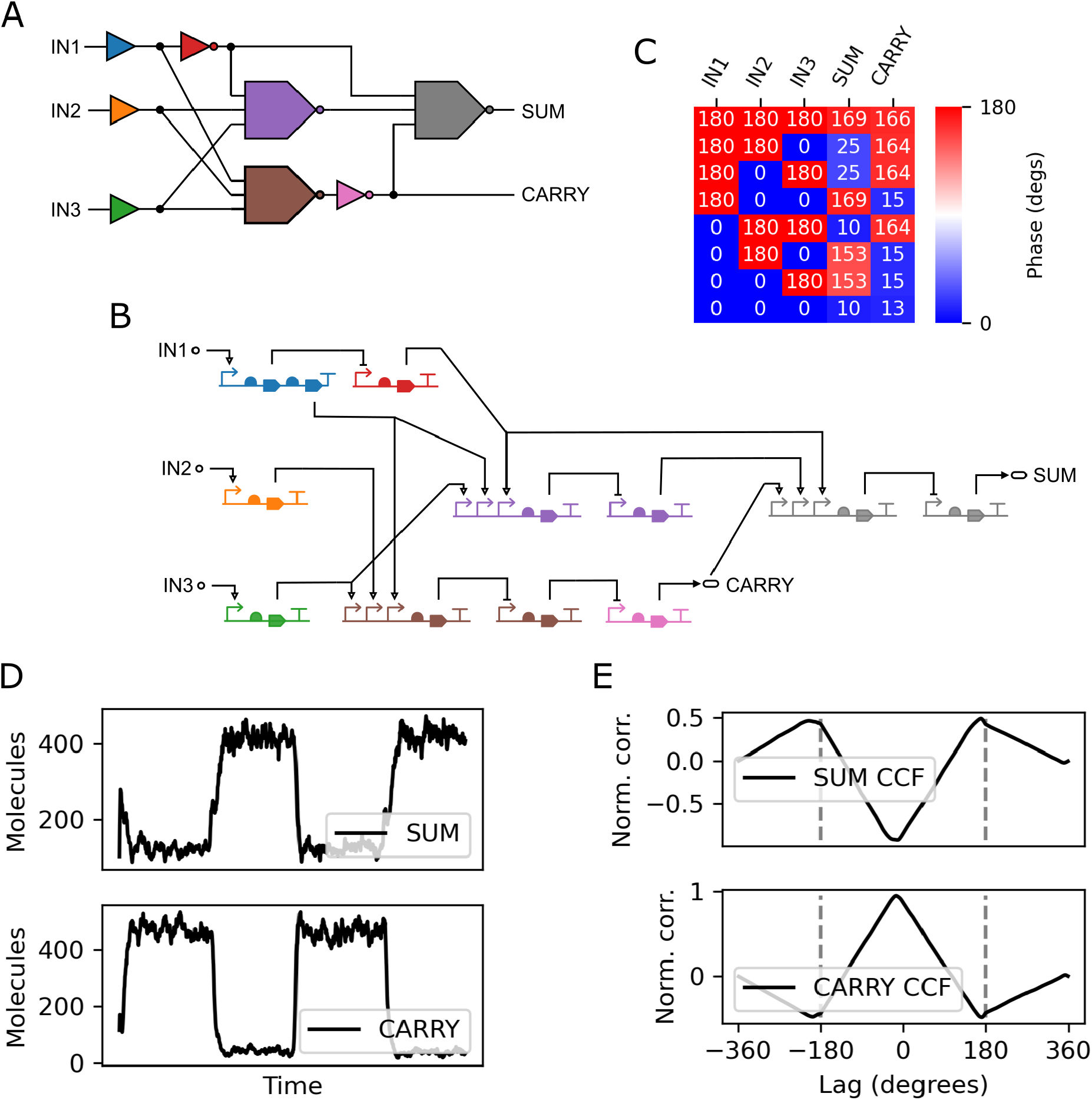
Design of phase-based genetic full adder circuit. (A) Schematic of the circuit with three input signal receivers representing the input bits to add (IN1 and IN2) and the carry bit (IN3). (B) Genetic implementation of the circuit in A. (C) Logical truth table showing mean phase of 100 stochastic simulations, where antiphase indicates logical 0 and in phase represents logical 1. (D) Example stochastic simulation traces for input normalized signal concentration and output protein level for computing the sum 1 + 1. (E) Normalized cross-correlation functions with respect to the reference signal for the examples in D.

Stochastic simulations with *α* = 500 (giving minimum switching point of around 50 molecules) and *δ* = 100 showed that the circuit was fully operational with zero error rate (figure 6C). Figure 6D shows an example computation with input IN1 = 0, IN2 = 1, and IN3 = 1. The correct logic is reproduced with SUM antiphase to the reference (logical 0), and CARRY in phase (logical 1). This can be seen from the cross-correlation of each protein level with the reference input signal (figure 6E).

Combining the full adder circuits in a cascade it is possible to perform addition of multi-bit numbers with a ripple adder circuit. We extended our full adder design to produce a 4-bit phase-based genetic ripple adder circuit (figure 7A). We performed 100 stochastic simulations of all 256 possible additions (*α* = 500, *δ* = 100), showing the circuit was fully functional with zero error rate. Figure 7B shows protein level traces for an example addition computation.

**Figure 7:**
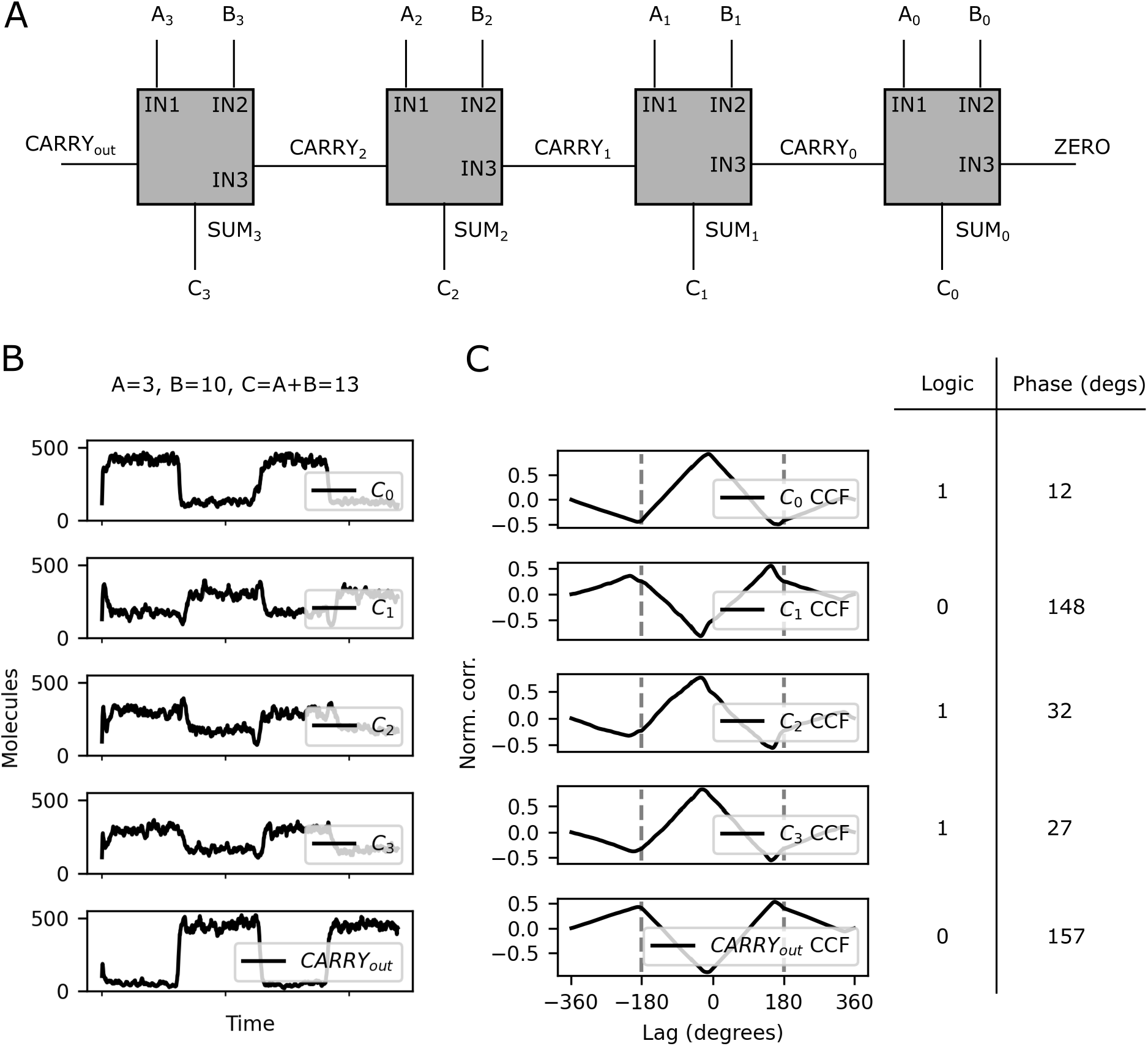
Design of a 4-bit phase-based genetic ripple adder circuit. (A) Schematic design of the circuit combining four full adders (gray boxes). (B) Example stochastic simulation traces for the computation *A* + *B* = 3 + 10 = 13. (C) Normalized cross-correlation functions (CCF) with respect to the reference signal for the traces in B. On the right are the phases and corresponding logic showing the correct outputs.

## Discussion

In this study we have designed and simulated genetic circuits that encode logical states in oscillating levels of gene expression. We showed that these circuits could perform computation that scales to multiple layers of sequential logic, demonstrated by the design of a 4-bit ripple adder. Furthermore, our results predict that these phase-based genetic logic circuits would be less sensitive to gate mismatch than level-based genetic circuits. This means that they would be more robust to inaccuracies in estimates of model parameters.

We showed that multiple complete logical sets can be formed using phase-based genetic logic. In level-based genetic logic, the NOR gate has been used to synthesise many logic circuits, including adders and digital displays (*4, 5, 15, 16*). However, these circuits may not be the most efficient. The use of complementary MAJORITY and NOT gates may enable more efficient design of some circuits, reducing the number of proteins expressed and thus metabolic burden on cells when compared to level-based NOR gate logic.

There are however some practical challenges to overcome in the implementation of the phase-based genetic logic designs presented here. Transcriptional interference between promoters in the MAJORITY and complementary MAJORITY gate designs (utilizing triple promoters) may hinder their correct function. Such interference has been demonstrated in constitutive promoters (*17*), but to our knowledge remains to be studied for regulated promoters as used in our designs. Similar to level-based logic, scaling to large circuits requires multiple orthogonal transcriptional activators and repressors. Large libraries of repressors have been created (*5*), but a limited number of transcriptional activators are currently available (*18*). Again similar to level-based logic, the number of gates in a circuit is limited by metabolic burden, which may be mitigated by splitting circuits into multiple communicating strains in consortia (*19, 20*).

Further development and application of the phase-based genetic logic approach will require logic synthesis to the range of gates designed in this study. Such tools might build on work in quantum computing which utilizes MAJORITY logic synthesis (*21, 22*). Building on logic synthesis we envisage full genetic design automation such as that implemented in the Cello (Cellular Logic) software (*5*), which compiles a required logical truth table into an output genetic design. This process utilizes a library of characterized gates, which could be developed for the designs presented here. Finally, a natural extension to this approach is to couple phase-based genetic logic gates to genetic oscillators (*13, 23–25*) to drive their inputs. In this case, sub-harmonic injection locking of genetic oscillators might also be exploited to store logical states, producing a finite state machine (*10*).

## Methods

All simulations were performed using the LOICA (Logical Operators for Integrated Cell Algorithms) Python package (*26*). Plots were generated using the Matplotlib Python package (*27*) and Jupyter notebooks (*28*).

## Acknowledgement

The authors thank Jaijeet Roychowdhury for helpful discussions. The authors are supported by the Newcastle University School of Computing. This research made use of the Rocket High Performance Computing service at Newcastle University.

